# Metaplastic neuronal state transition regulates species-specific interoceptive processing in *Drosophila*

**DOI:** 10.64898/2026.02.07.702027

**Authors:** Adam M. Brann, Dieu Linh Nguyen, Alexa N. Zarjetskiy, Natalia Pokaleva, Elizabeth M. Paul, Masashi Tabuchi

**Author notes:** These authors have contributed equally to this work (Adam M. Brann, Dieu Linh Nguyen, Alexa N. Zarjetskiy).

## Abstract

Interoceptive processing, which involves the sensing and integration of internal physiological states, is fundamental to maintaining homeostasis. However, how species-specific interoceptive features arise from the underlying biophysical properties of neural circuit physiology remains unclear. We investigate the biophysical basis of species-specific interoception by examining protein-hunger dopamine neurons (DA-WED) in two *Drosophila* species with divergent dietary ecologies. We find that DA-WED neurons in *D. melanogaster* exhibit weak persistence of internal states, enabling flexible behavioral transitions during nutrient stress. In contrast, *D. sechellia* shows strong state persistence, locking neurons into a “preferred” configuration during protein deprivation. This divergence is supported by distinct intrinsic membrane properties, including protein deprivation-induced rebound spikes unique to *D. sechellia*. Analysis of synaptic dynamics and cardiomyocyte electrophysiology reveals species-specific physiological regulations coordinating central and peripheral systems. Behavioral assays confirm corresponding differences in protein consumption strategies, directly linking neural state geometry to ecologically relevant feeding behavior. Our findings establish metaplastic regulation of neural state transitions as a fundamental mechanism through which ecological specialization shapes interoceptive processing and brain-body coordination.

**Significance Statement:** We identify physiological regulations of neural state transitions as a core mechanism underlying species-specific interoceptive processing. Through comparative electrophysiology in *D. melanogaster* and *D. sechellia*, we demonstrate that ecological specialization manifests through distinct intrinsic membrane properties of DA-WED neurons, fundamentally altering neural state space geometry during protein deprivation. Species-specific synaptic plasticity gates these transitions while coupling cardiac rhythms to central computation. Our findings reveal how evolution transforms nutrient sensing into divergent neural dynamics, establishing a mechanistic framework for understanding how ecological pressures sculpt the biophysical architecture of interoceptive circuits to coordinate adaptive brain-body interactions.

## Introduction

The ability of the brain to detect, integrate and act upon internal physiological states, known as interoceptive processing (1), is essential for maintaining homeostasis in fluctuating environmental conditions (2, 3). Interoception is not a static readout of peripheral signals but a dynamic process in which internal bodily rhythms, including metabolic, autonomic, and cardiac signals, are centrally integrated to shape neural computation and behavior (4-7). While significant progress has illuminated interoceptive mechanisms within individual species, a critical gap remains in understanding how evolution shapes interoceptive processing to match species-specific ecological demands and what neural circuit mechanisms underlie this specialization. Homeostasis depends not only on preserving internal stability (8, 9) but also on the capacity for flexible behavioral adaptation when internal resources are constrained (10). Feeding decisions provide a powerful example of this principle (11-14). Rather than responding solely to caloric deficits, animals exhibit nutrient-specific appetites, engaging dedicated neural circuits that detect deficiencies in particular macronutrients such as protein or lipid (15-17). These behaviors emerge from the integration of internal physiological signals with central neural processing, transforming interoceptive inputs into adaptive motor and autonomic outputs. Emerging evidence suggests that species-specific behavioral differences cannot be explained solely by variation in peripheral sensory detection but instead reflect divergence in how internal signals are transformed within central neural circuits (18). Evolutionary studies of taste processing, for example, demonstrate that dietary specialization can be driven by changes in central sensorimotor transformations rather than peripheral receptor repertoires (19-22). This raises the possibility that species-specific interoception may likewise emerge from differences in neural state dynamics and plasticity, rather than from differences in sensory detection alone. Protein hunger neurons in *Drosophila* provide an ideal system for investigating these questions. DA-WED neurons in *Drosophila* were characterized as a single class of dopamine neurons representing protein hunger with their membrane excitability and synaptic plasticity (23), and defining setpoint based on their resting membrane potential (24). However, the ways in which ecological pressures potentially shape the neural basis of nutrient homeostasis and the properties of protein hunger representation across species with different dietary strategies remain largely unknown. We tackled this question using comparative neurophysiological approaches by using DA-WED neurons in *D. melanogaster* and *D. sechellia*. These closely related species exhibit dramatically different protein requirements and feeding strategies despite sharing recent evolutionary history (21, 25-31). We hypothesized that their behavioral differences arise not from altered sensory capabilities but from fundamental divergence in neural state dynamics and plasticity mechanisms. Our results support this hypothesis, revealing that protein deprivation triggers profoundly different patterns of activity of DA-WED neurons with distinctive intrinsic membrane properties and synaptic inputs, and brain-heart coupling between species. These findings establish metaplastic modulation of neural state transitions as a central mechanism linking ecological specialization to interoceptive processing, demonstrating how evolution reprograms the computational geometry of neural circuits to generate species-appropriate homeostatic responses.

## Results

### Distinguished intrinsic membrane properties of DA-WED neurons in *D. sechellia* induced by protein deprivation

To examine species-specific neural responses to protein deprivation, we compared the intrinsic membrane properties of DA-WED neurons in *D. melanogaster* and *D. sechellia*, hypothesizing that protein deprivation induces distinct electrophysiological properties in *D. sechellia*. Using intracellular recordings combined with the retrograde labeling technique, we examined intrinsic membrane properties of DA-WED neurons to compare them between two *Drosophila* species with and without protein deprivations (**Fig. 1*A***). We found that *D. sechellia* DA-WED neurons exhibited significantly higher mean firing rates in their spontaneous firing activity when they are exposed to protein deprivation, compared to those in *D. melanogaster* (**Fig. 1*B***), while we found no difference in temporal structural analysis based on inter-spike interval distribution by using spike irregularity of global (**Fig. 1*C***) and local metrics (**Fig. 1 *D* and *E***). To assess stochastic state transition in spike sequence structures, we constructed a discrete-time Markov chain model based on action potentials of DA-WED neurons (**Fig. 1*F***). Action potential waveforms were classified into three discrete states based on a k-means clustering algorithm-based sorting. We found that Markov chain analysis revealed that DA-WED neurons in *D. sechellia* exhibited stronger state persistence compared to *D. melanogaster*, which was further strengthened following protein deprivation. To investigate the biophysical basis of a state-transition probability model derived from spike waveform–based clustering, we analyzed action potential waveforms and found that the temporal kinetics of afterhyperpolarization (AHP) are significantly altered in DA-WED neurons of *D. sechellia* following protein deprivation (**Fig. 1 *G, H* and *I***), potentially suggesting species-specific divergence in ion channel properties involved in forming AHP that was sensitive to protein deprivation. To determine whether protein deprivation selectively alters specific aspects of the intrinsic excitability of DA-WED neurons, we examined their responses to hyperpolarizing current injections in both *D. melanogaster* and *D. sechellia* (**Fig. 1*J***). Strikingly, DA-WED neurons in protein-deprived *D. sechellia* generated rebound spikes when hyperpolarizing stimuli were terminated. This rebound activity was not observed under any other conditions, demonstrating that rebound spiking emerges specifically in *D. sechellia* under protein-limited conditions (**Fig. 1*K***). These findings suggest that the acquisition of rebound excitability in DA-WED neurons is species-specific and nutritionally gated, rather than being a general property of these neurons or a universal response to protein deprivation. Such conditional rebound firing may enable the rapid reactivation of DA-WED neurons following inhibition, potentially representing an adaptive mechanism that supports protein-seeking behaviors in the protein-restricted ecological niche of *D. sechellia*. We did not find any significant differences in other functional parameters related to intrinsic membrane properties (**Fig. *S1***).

**Fig. 1:**
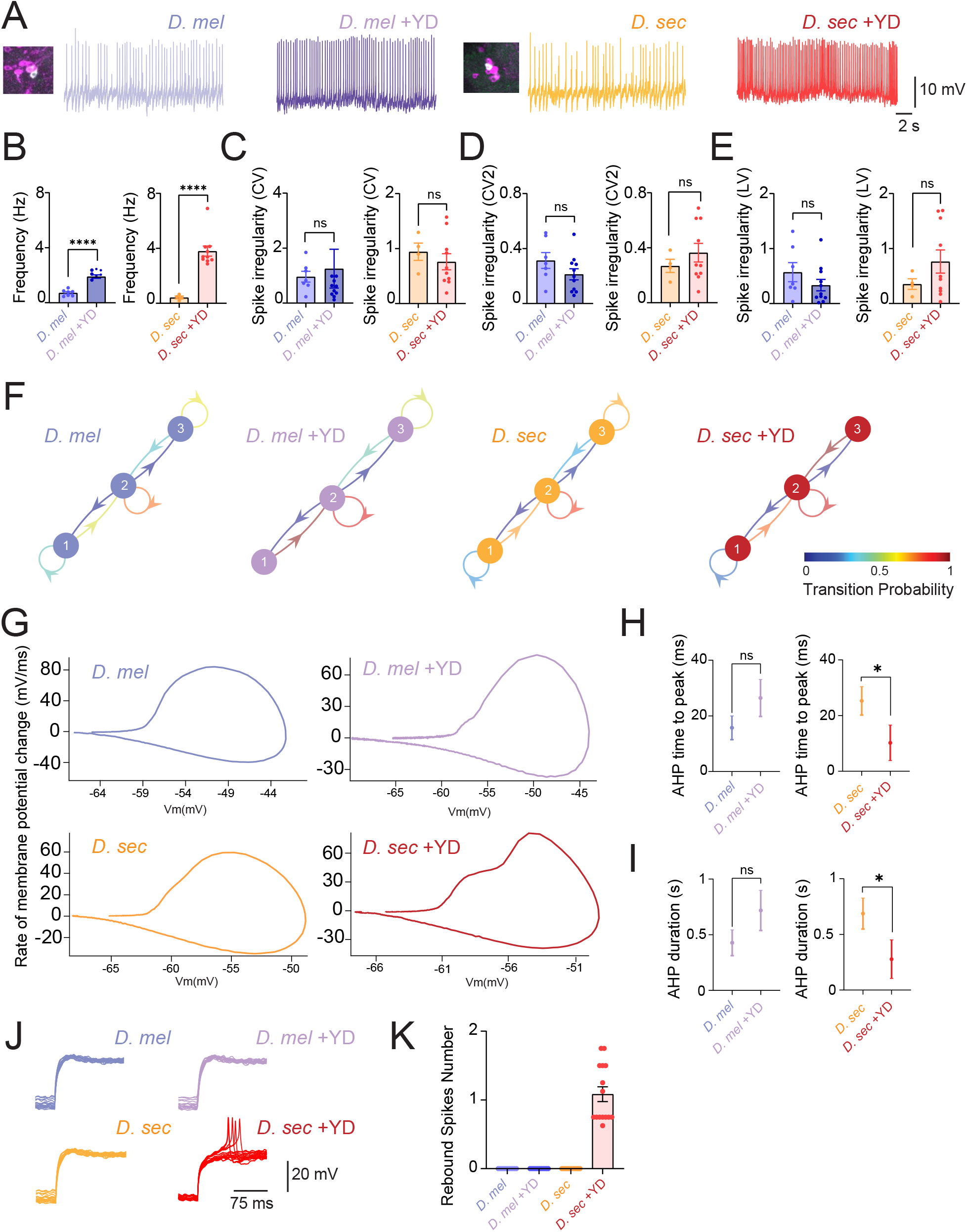
Distinguished intrinsic membrane properties of DA-WED neurons in *D. melanogaster* and *D. sechellia*. (**A**) Membrane voltage traces in spontaneous firing activity in DA-WED neurons in *D. melanogaster* and *D. sechellia* with and without protein deprivation. Cell body imaging with retrograde labeling technique to identify DA-WED neurons in *D. melanogaster and D. sechellia* (**B**) Quantification of mean firing rate. (**C**) Quantification of spike time irregularity using the coefficient of variation (CV) (**D**) Quantification of spike time irregularity using the local variation coefficient (CV_2_) (**E**) Quantification of Spike time irregularity using the local variation (LV) metrics (**F**) Stochastic process of state transition visualized by discrete-time Markov chains in DA-WED neurons with and without protein deprivation in *D. melanogaster* and *D. sechellia*. Categorization of the quantified maximum spike onset velocity (dV/dt) into three different classes based on k-means clustering. The figure shows the transition matrix generated to calculate the probability of transitioning between these defined classes. (**G**) Phase plot representation of action potential waveform, with membrane potential plotted against its rate of change (dv/dt). Trajectories reflect the instantaneous state of the membrane potential dynamics neuron action potential generation. (**H**) Quantification of the interval between the action potential peak and the minimum membrane potential reached during the subsequent afterhyperpolarization (AHP). (**I**) Quantification of the duration of AHP, defined as time during which membrane potential remained below baseline following an action potential (**J**) Membrane voltage traces showing rebound spiking within DA-WED neurons following current injections (**K**) Quantification of the amount of rebound spiking seen in each of the four groups, only being seen in *D. Sechellia* during protein deprivation. Sample sizes: *D. melanogaster*: n=7, *D. melanogaster* with protein deprivation: n=11, *D. sechellia*: n=4, *D. sechellia* with protein deprivation: n=10 *p < 0.05, ****p < 0.0001. ns: non-significance based on unpaired t-tests.

### Distinguished synaptic inputs in DA-WED neurons associated with dynamical stochastic process

To investigate whether species-specific differences in DA-WED neurons can be found in their synaptic input properties under protein deprivation, we compared the synaptic properties of DA-WED neurons between *D. melanogaster* and *D. sechellia*. We hypothesized that the heightened sensitivity to protein deprivation observed in *D. sechellia* could be explained by distinctive synaptic input features in DA-WED neurons. Using intracellular recordings, we quantified postsynaptic potentials (PSPs) in DA-WED neurons and found that PSP amplitudes were significantly larger in *D. sechellia* than in *D. melanogaster* (**Fig. 2*A***), and these greater PSP amplitudes in *D. sechellia* were further enhanced following protein deprivation (**Fig. 2*B***). We next analyzed PSP frequency based on their inter PSP event intervals and instantaneous frequency (**Fig. *S2 A and B***) and found that *D. sechellia* exhibited a higher PSP frequency than *D. melanogaster*, which was further increased under protein-deprived conditions (**Fig. 2 *C* and *D***). To assess whether the temporal dynamics of synaptic inputs were also altered, we applied a Gaussian Mixture Model (GMM) to characterize the structure of PSP timing. The inter event intervals of two adjacent PSPs were projected into a two-dimensional space, and a GMM was used to estimate the geometry of their joint probability distribution (**Fig. 2*E***). Under control conditions in both species, PSP timing was best described by a single cluster, characterized by a reduced probability of tail events and no discrete separation in stochastic distance. In contrast, protein deprivation induced the emergence of additional clusters, with distinct species-specific patterns. Protein-deprived *D. melanogaster* exhibited multiple large, well-separated clusters, whereas protein-deprived *D. sechellia* displayed multiple smaller, fragmented clusters. To investigate how biophysical changes in intrinsic membrane properties relate to altered synaptic dynamics, we implemented computational modeling of DA-WED neuron activity. We hypothesized that hidden temporal structures in synaptic potentials might not be immediately apparent from raw recordings. To test this, we modeled the time series of membrane potentials using an Ornstein–Uhlenbeck (OU) process and analyzed the dynamics with continuous wavelet transforms (**Fig. 2*F***). By using OU parameters, including the time constant (**Fig. *S2C***) and noise amplitude (**Fig. *S2D***), we identified dramatic differences in the temporal structure of continuous wavelet transform dynamics between DA-WED neurons under protein deprivation and controls (**Fig. 2 *G* and *H***). The variability was significantly larger in DA-WED neurons experienced protein deprivation. Together, these results support DA-WED synaptic plasticity as a metaplastic regulator of neural state transitions underlying species-specific protein hunger neurophysiological reaction in two *Drosophila* species.

**Fig. 2:**
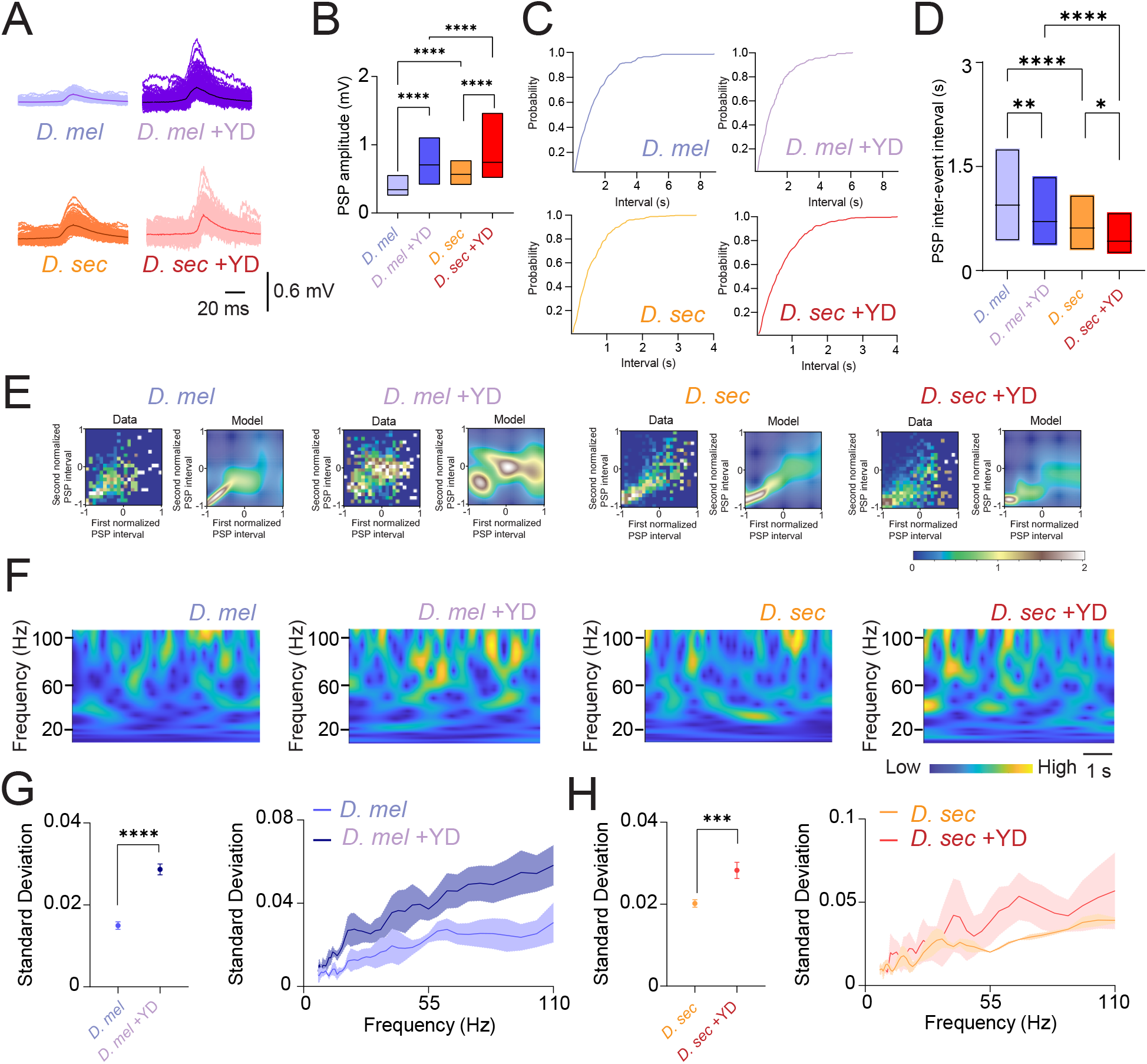
Distinguished synaptic inputs in DA-WED neurons in *D. melanogaster* and *D. sechellia*. (**A**) Superimposed membrane potential traces for spontaneous PSPs of DA WED neurons in *D. melanogaster* and *D. sechellia* with and without protein deprivation (**B**) Quantification of PSP amplitudes (**C)** Cumulative probability of PSP in *D. melanogaster* and *D. sechellia* with and without protein deprivation (**D**) Quantification of inter-event intervals of the PSP distribution (**E**) The interval times of two adjacent PSPs were projected into two-dimensional space, and a Gaussian Mixture Model (GMM) was used to computationally capture the geometry of their probability distribution. Sample sizes: *D. melanogaster*: n=4, *D. melanogaster* with protein deprivation: n=7, *D. sechellia*: n=3, *D. sechellia* with protein deprivation: n=8. *p < 0.05, **p < 0.01, and ****p < 0.0001 based on one-way ANOVA followed by post-hoc Tukey tests. (**F**) Continuous wavelet transform in the time series of the Ornstein–Uhlenbeck (OU) model obtained from membrane voltage traces in DA-WED neurons. (**G**) Quantification of the differences in Standard Deviation of the OU model within *D. melanogaster* with and without protein deprivation (**H**) Quantification of the differences in Standard Deviation of the OU model within *D. sechellia* with and without protein deprivation. Sample sizes: *D. melanogaster*: n=3, *D. melanogaster* with protein deprivation: n=6, *D. sechellia*: n=3, *D. sechellia* with protein deprivation: n=3. ***p < 0.001 based on unpaired t-tests.

### Distinguished electrophysiological properties of cardiomyocytes

Interoceptive processing coordinates internal physiological states through interorgan communication, with the brain–heart axis serving as a key mechanism by which organisms maintain internal stability in response to environmental change (32-34). To examine whether the species-specific electrophysiological signatures observed in DA-WED neurons extend beyond the central nervous system, we performed intracellular recordings from cardiomyocytes in *D. melanogaster* and *D. sechellia* under protein-deprived conditions (**Fig. 3 *A* and *B***), and analyzed their functional parameters, including inter-spike interval distribution (**Fig. *S3***). We found that both *D. melanogaster* and *D. sechellia* cardiomyocytes showed a trend toward increased mean firing rates under protein deprivation compared with baseline controls, although these differences did not reach statistical significance (**Fig. 3 *C***). Analysis of the temporal structure of cardiac action potentials revealed pronounced interspecies differences in firing variability. Interestingly, we found that global variability metrics indicated increased irregularity following protein deprivation (**Fig. 3 *D***), whereas local variability metrics revealed reduced irregularity in *D. sechellia* cardiomyocytes, as measured by CV2 and LV (**Fig. 3 *E* and *F***).

**Figure 3.**
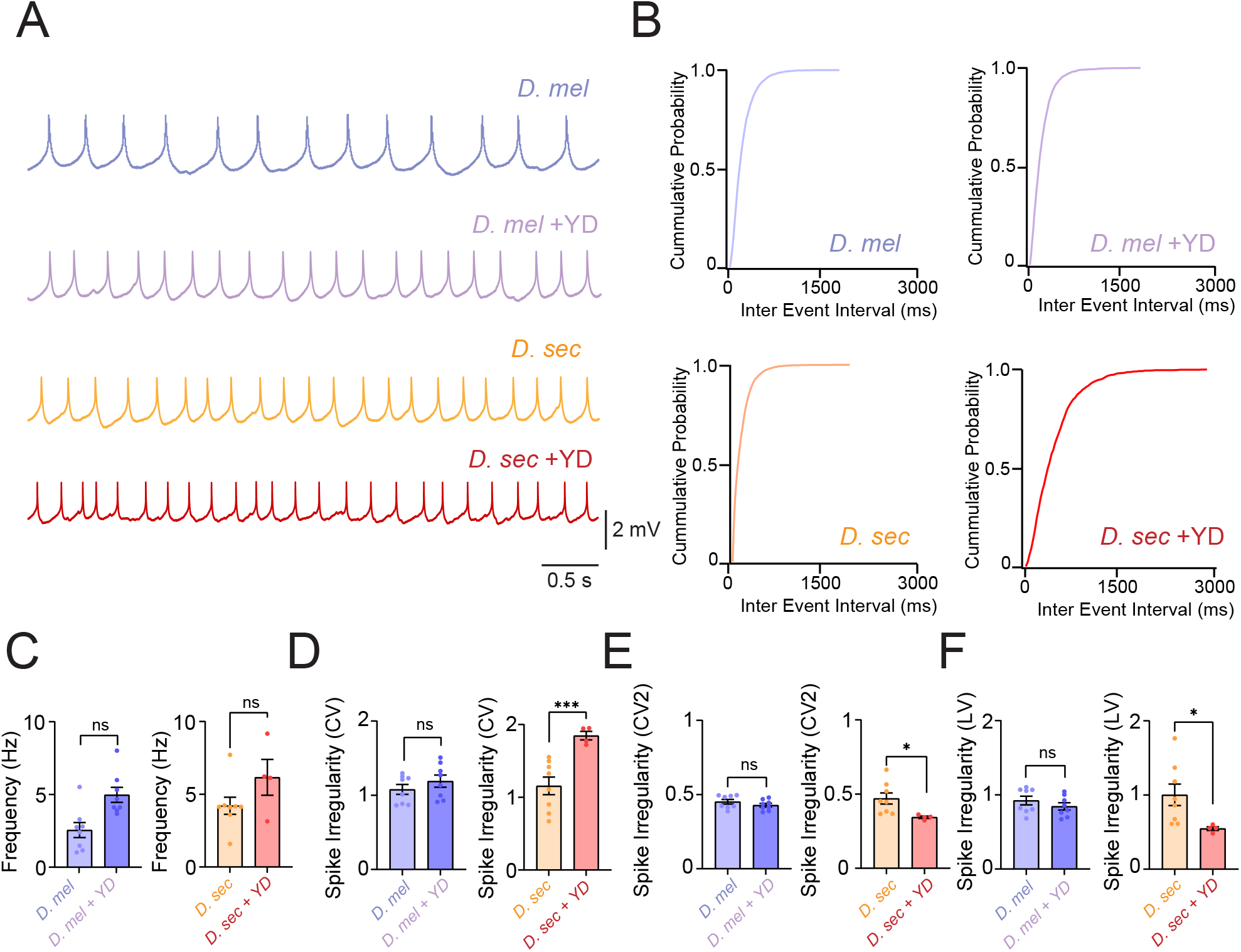
Distinguished cardiomyocytes activity in *D. melanogaster* and *D. sechellia*. (**A**) Representative membrane voltage traces illustrating spontaneous activity patterns of cardiomyocytes under control and yeast-deprived conditions in *D. melanogaster* and *D. sechellia*. (**B**) Cumulative probability of inter event interval distribution of spontaneous activity patterns of cardiomyocytes under control and yeast-deprived conditions in *D. melanogaster* and *D. sechellia*. (**C**) Quantification of mean activity rates of spontaneous activity of cardiomyocytes under control and yeast-deprived conditions in *D. melanogaster* and *D. sechellia*. (**D**) Quantification of irregularity based on CV, reflecting global variability of inter event interval distribution of spontaneous activity patterns of cardiomyocytes under control and yeast-deprived conditions in *D. melanogaster* and *D. sechellia*. (**E**) Quantification of irregularity (CV2), reflecting local variability of inter event interval distribution of spontaneous activity patterns of cardiomyocytes under control and yeast-deprived conditions in *D. melanogaster* and *D. sechellia*. (**F**) Quantification of spike timing irregularity (LV), reflecting local inter event interval distribution of spontaneous activity patterns of cardiomyocytes under control and yeast-deprived conditions in *D. melanogaster* and *D. sechellia*. Sample sizes: *D. melanogaster*: n=8, *D. melanogaster* with protein deprivation: n=8, *D. sechellia*: n=8, *D. sechellia* with protein deprivation: n=4. *p < 0.05, and ***p < 0.001. ns: non-significance based on unpaired t-tests.

**Figure 4.**
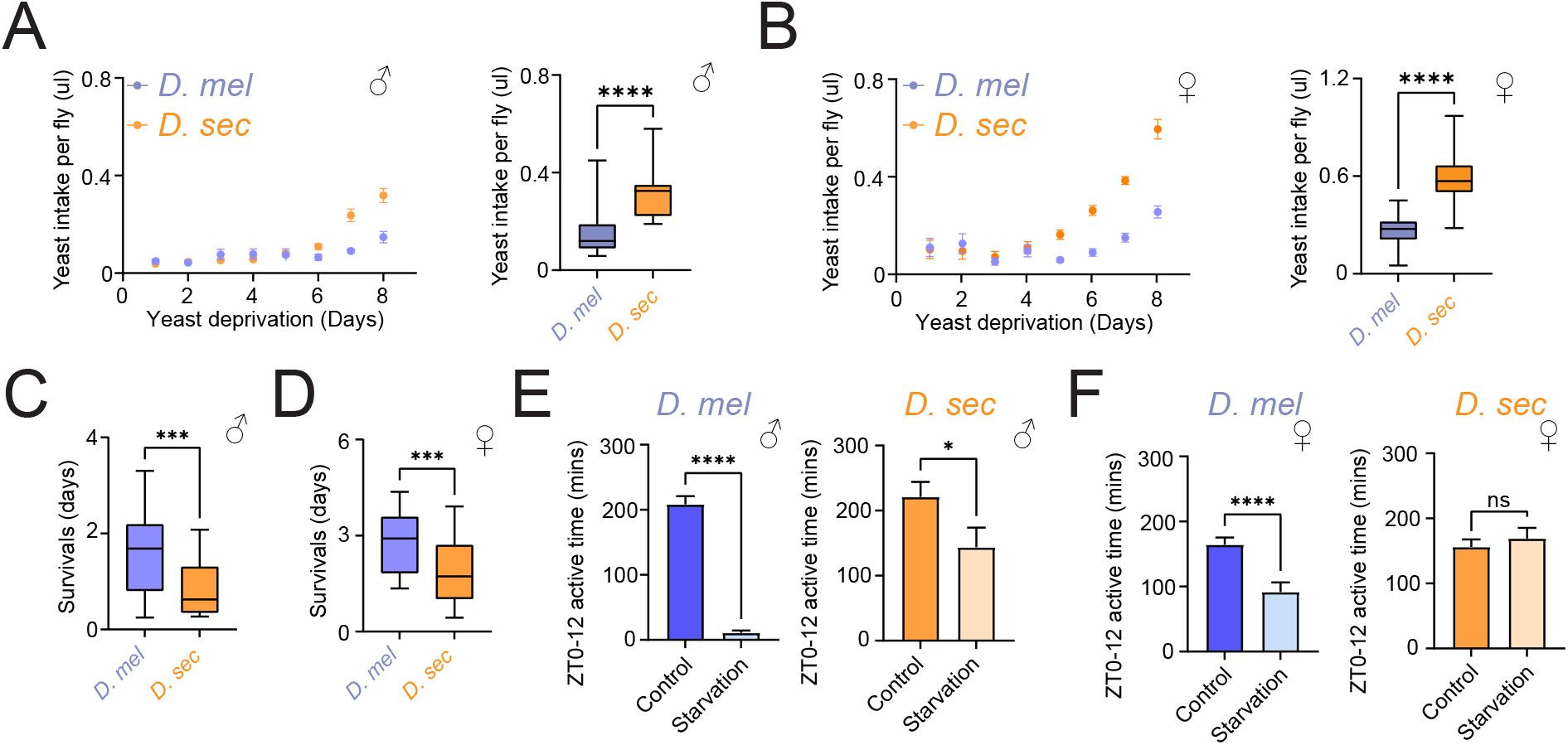
Yeast intake, survival and activity rates during starvation periods. (**A-B**) Quantification of yeast intake per fly (ul) after yeast deprivation (days) in males (**A**) and females (**B**). Sample sizes: *D. melanogaster* (male): n=16, *D. sechellia* (male): n=16, *D. melanogaster* (female): n=16, *D. sechellia* (female): n=16. ****p < 0.0001 based on unpaired t-tests. (**C-D**) Comparison of survival length (days) during starvation periods in males (**C**) and females (**D**). Sample sizes: *D. melanogaster* (male): n=29, *D. sechellia* (male): n=28, *D. melanogaster* (female): n=32, *D. sechellia* (female): n=32. ****p < 0.0001 based on unpaired t-tests. (**E**) Comparison of daytime (ZT0-12) active time (mins) under control and starvation conditions in males. (**F**) Comparison of daytime (ZT0-12) active time (mins) under control and starvation conditions in females. Sample sizes: *D. melanogaster* (male): n=160, *D. melanogaster* (male) with starvation: n=160 *D. sechellia* (male): n=54, *D. sechellia* (male) with starvation: n=64, *D. melanogaster* (female): n=160, *D. melanogaster* (female) with starvation: n=160, *D. sechellia* (female): n=64,, *D. sechellia* (female) with starvation: n=64. *p < 0.05, and ***p < 0.001, and ****p < 0.0001. ns: non-significance based on unpaired t-tests.

### Distinguished feeding behaviors between *D. melanogaster* and *D. sechellia*

Given the distinct dietary ecologies of *D. melanogaster* and *D. sechellia* (35-37), we hypothesized that evolutionary divergence in the biophysical properties of DA-WED neurons encoding protein hunger would manifest not only in neural state dynamics, but also in behavioral phenotypes related to nutrient intake, energy balance and activity. Thus, we performed feeding behavioral assays to determine how neural dynamics scale to whole-animal physiology and behavior. We found that in *D. sechellia*, protein intake did not differ from baseline during the first five days of early adulthood in either sex. However, a significant increase in protein intake was observed during the final three days of early adulthood in both males (**Fig. 5 *A***) and females (**Fig. 5 *B***). Despite this increased protein consumption, *D. sechellia* displayed reduced starvation tolerance compared to the control groups, and this reduction was evident in both males (**Fig. 5 *C***) and females (**Fig. 5 *D***). There were marked species-specific differences in daytime locomotor activity. In *D. melanogaster*, the amount of time spent active during the day was significantly reduced in both males (**Fig. 5 *E***) and females (**Fig. 5 *F***). In contrast, *D. sechellia* did not exhibit a comparable reduction in daytime activity. Male *D. sechellia* showed only a minor decrease in active time during the day compared to *D. melanogaster* males, while female *D. sechellia* showed no reduction compared *to D. melanogaster* females.

## Discussion

Our comparative neurophysiological analyses show that DA-WED neurons in *D. melanogaster* exhibit weak persistence of internal neural states, whereas neurons in *D. sechellia* display strong bias toward a single “preferred” state under protein deprivation. Protein deprivation not only reprograms the intrinsic dynamics of DA-WED neurons but also alters synaptic inputs and cardiomyocyte electrophysiology, indicating coordinated plasticity across multiple organs. These state-dependent dynamics are likely supported by distinct intrinsic membrane properties, including the emergence of rebound spiking uniquely in *D. sechellia* under protein deprivation. Together, these findings highlight the role of interoceptive rhythms and brain–body coordination in shaping adaptive responses to metabolic stress. Building on these observations, we propose that metaplastic regulation of neural state transitions may represent a general evolutionary strategy for optimizing interoceptive processing. By tuning intrinsic excitability and synaptic plasticity, neural circuits can reprogram their state space to meet species-specific ecological demands while preserving homeostatic stability. More broadly, this framework provides mechanistic insight into how neural circuits evolve to integrate internal physiological signals with adaptive behavior, revealing principles governing coordinated brain–body regulation across species. Our electrophysiological characterization demonstrated that *D. sechellia* DA-WED neurons exhibited enhanced responsiveness to protein deprivation through multiple coordinated mechanisms, including increased spontaneous firing rates and distinctive intrinsic properties, such as altered afterhyperpolarization kinetics and rebound spiking. Analyses of neural state-space geometry indicate that these intrinsic features are critical determinants of species-specific responses to protein deprivation, possibly enabling *D. sechellia* to rapidly reprogram feeding behavior in response to protein availability. Global (CV) and local (CV2 and LV) variability metrics initially appeared contradictory due to the temporal scales they capture: CV reflects long-term fluctuations, while CV2 and LV measure local (38, 39). In our study, protein deprivation increased CV but reduced CV2 and LV, indicating that cardiomyocytes’ action potential timing remained stable while broader temporal irregularity grew. These results suggest that interoceptive challenges engage multiscale mechanisms along the brain–heart axis, reflecting coordinated regulation between central and peripheral tissues. This coordination is consistent with our observation that cardiac electrophysiological dynamics mirror activity patterns observed in DA-WED neurons, indicating species-specific coupling between neural state regulation and peripheral physiology. From an ecological perspective, such adaptations may confer selective advantages to *D. sechellia*. Although this species exploits protein-poor morinda fruit as its primary resource (25), reproductive and developmental demands likely necessitate periodic episodes of intensive protein intake. The delayed yet intensified protein drive observed in *D. sechellia* may therefore optimize energy utilization under chronic protein scarcity, while preserving the capacity for rapid protein acquisition when resources become available. This biphasic interoceptive response aligns with the temporal structure of protein availability in *D. sechellia*’s natural environment, where morinda fruit ripening cycles generate transient windows for supplemental protein intake. We also showed that *D. melanogaster* showed decreased daytime activity during prolonged, lethal starvation periods as compared to *D. sechellia* which showed little change. Food-seeking behaviors are known to be dopamine-modulated in *Drosophila* (40). A maintenance of locomotor activity in *D. sechellia* might be due to elevated baseline levels of DA caused by the Catsup mutation (41).The ecological specialization of *D. sechellia* is driven by a need to regulate L-DOPA levels through dietary intake of the noni fruit (41). We propose that this combination of elevated baseline dopamine and diet-dependent l-DOPA supply allows DA levels in *D. sechellia* to remain relatively high even during starvation, thereby continuing to promote foraging behaviors and sustaining locomotor activity. In *D*.*melanogaster*, in contrast, increased sleep and reduced activity under food deprivation are associated with enhanced starvation resistance (42, 43), consistent with the sharp drop in daytime locomotion and longer survival that we observed, suggesting a unique phenotypic difference between specialist and generalist species. The *D. sechellia* strategy, which enhances performance in its ecological niche, is accompanied by higher activity and lower survival under complete starvation. This may suggest that adaptations critical for ecological specialization may reduce fitness in other environmental contexts. Together, these findings show that dietary specialization entails fundamental reprogramming of interoceptive neural computation, spanning intrinsic neuronal properties to brain–body coordination, and involves trade-offs between ecological efficiency and physiological flexibility.

### Limitations of the study

Several factors limit the generalizability of our findings. First, DA-WED neurons in *D. sechellia* were identified by retrograde labeling followed by post hoc TH immunostaining, defining recorded cells as TH-positive neurons projecting to the WED neuropil. Although this approach is less rigorous than genetic labeling available in *D. melanogaster*, it provides anatomically grounded identification and does not undermine the comparative electrophysiological conclusions of this study. Second, we showed a correlation between DA-WED neuronal and cardiomyocyte electrophysiological phenotypes, but does not establish causality. While causal relationships will require future genetic perturbation tools in *D. sechellia*, the observed correlations still support the central claims of species-specific neurophysiological differences. Third, we focused solely on protein deprivation, although natural environments involve dynamically fluctuating nutritional conditions across multiple macro- and micronutrients, and DA-WED neurons represent only one node within a broader interoceptive network, including mushroom bodies (40, 44) and insulin-producing cells (45-48), should contribute to species-specific feeding behaviors through mechanisms we have not examined.

## Materials and Methods

### *Drosophila* genetics

*D. melanogaster* used in this study was generated by crossing *TH-D-Gal4* with *TH-C-FLP*; *UAS-FRT-STOP-FRT-mCD8-GFP* to drive expression of *mCD8-GFP* where both Gal4 and FLP were active (*TH-D-Gal4, TH-C-FLP>UAS-FRT-STOP-FRT-mCD8-GFP*), allowing DA-WED neurons to visualize under with GFP fluorescence (23, 24). These fly lines were gifts from Mark Wu. Flies were kept in an incubator (DR-36VL, Percival Scientific, Perry, IA, United States) under 12 h:12 h light-dark cycles and 65% humidity. Files were fed standard *Drosophila* food containing molasses, cornmeal, and yeast. *D. sechellia* strain K-S10 (49) was obtained from KYORIN-Fly (Fly Stocks of Kyorin University, Japan) and was maintained in a standard *Drosophila* food having Noni fruit sprinkles. Flies used for all experiments were female with the age range of 5-8 days old. All experiments were conducted at 25 °C and were performed in compliance with all relevant ethical regulations for animal testing and research at Case Western Reserve University.

### Electrophysiology

We conducted electrophysiology by using sharp electrode intracellular recordings as described in previous studies (50-52), but recording stability challenges associated with conventional electrodes prompted adoption of membrane-coating technology described previously (53, 54). This approach required synthesizing lipid coatings from egg-derived lecithin (440154 from Sigma-Aldrich, concentration 7.6 g/L) combined with cholesterol (C8667 from Sigma-Aldrich, concentration 2 mM). These components were dissolved by ultrasonication for 60 min in an organic solvent mixture of hexane and acetone at room temperature. Solvents were subsequently removed by nitrogen gas evaporation followed by vacuum desiccation. Subsequently, the lipid mixture was dissolved in paraffin-squalene blend (ratio 7:3 by volume) and incubated at 80 °C overnight for stabilization. Sharp microelectrodes (120-190 MΩ) were created from quartz glass capillaries measuring 1.2 mm outer and 0.6 mm inner diameter using laser-based puller (P-2000, Sutter Instrument). The electrodes were coated with phospholipids using a tip-dip technique (53), involving sequential exposure to internal solution and lipid layer with subsequent vertical withdrawal under micromanipulator guidance. Sharp electrode recordings employed 1 M KCl with as internal electrode solution with 13 mM biocytin hydrazide (B1603, ThermoFisher). All solutions underwent sterilization using 0.02 micrometer pore filters (Anotop 10, Whatman). To make electrophysiological preparations, flies underwent cooling-induced anesthesia before attachment to metal sheets using dental wax or ultraviolet-cured adhesives. Surgical access involved removing cuticular structures to reveal brain surfaces. Preparations were maintained in *Drosophila* saline containing: 101 mM NaCl, 3 mM KCl, 1 mM CaCl_2_, 4 mM MgCl_2_, 1.25mM NaH_2_PO_4_, 20.7 mM NaHCO_3_, and 5 mM glucose; with osmolarity adjusted to 235-245 mOsm and pH 7.2), which was pre-bubbled with 95% O_2_ and 5% CO_2_. Tracheal structures and muscle tissues were surgically removed. Glial barriers surrounding target neurons underwent enzymatic digestion using the treatment of the enzymatic cocktail, collagenase (0.1 mg/mL), protease XIV (0.2 mg/mL), and dispase (0.3 mg/mL), at 22°C for 1-2 min. Cardiac electrophysiological recordings were performed in dissected flies while preserving the structural integrity of the cardiac tube. This preparation maintained cardiac structure throughout the recordings, enabling reliable assessment of intrinsic cardiomyocyte properties using sharp microelectrode techniques. Intracellular recordings of cardiac electrical activity were obtained from cardiomyocytes using sharp microelectrodes. Electrodes used were the same as those used in DA-WED neurons but were filled with 3 M KCl and mounted on a flexible support wire to allow tracking of cardiac motion during recordings. Electrode insertion was monitored by changes in membrane potential and auditory feedback (Model 3300 Audio Monitor, A-M Systems). Membrane potentials were recorded once stable, typically several minutes after impalement. The difference between the intracellular electrode and an extracellular Ag/AgCl reference electrode was amplified using a high-impedance amplifier (Axoclamp 2B with HS-2A x1 LU headstage, Molecular Devices) and digitized via a Digidata 1550B interface at 10 kHz. Signals were low-pass filtered at 1 kHz, and data acquisition and analysis were performed using pCLAMP 11 (Molecular Devices) and Clampfit 10. Because cardiac muscle lacks classical neuromuscular junctions, the recorded depolarisations reflect cardiac action potentials or compound tissue responses, characterized by slow diastolic depolarisation followed by a rapid upstroke typical of invertebrate pacemaker activity (55, 56).

Local PSP variability was visualized using a Gaussian Mixture Model (GMM) following previous methods (52, 54). The GMM models data as a mixture of Gaussian components, each defined by a mean and covariance matrix, with mixture weights and component parameters estimated via the Expectation-Maximization (EM) algorithm. PSP interval timings were normalized to each cell’s average firing rate and aggregated across cells. Second-order distributions were constructed by logarithmically binning consecutive interval pairs into a 22 × 22 2D histogram, with logarithmic ranges adjusted for each experimental condition. Three to five full-covariance Gaussian components were fitted to these joint distributions. For validation, size-matched random samples generated from the GMM reproduced both joint and marginal distributions of the original data. To produce rate-matched PSP trains, an average PSP frequency was first sampled from a distribution fitted to PSPs. New trains were iteratively generated by sampling the next interval conditioned on the current interval using rejection sampling of the continuous conditional densities, after discarding 200 burn-in samples. The logarithmic intervals were exponentiated, normalized to PSP times, and converted into 10 ms binary signals.

Parameters extracted from membrane voltage recordings were used to construct an Ornstein–Uhlenbeck (OU) model. Membrane signals were preprocessed and mean-centered prior to analysis. The OU process’s time constant was derived from the temporal autocorrelation by fitting an exponential decay, equivalent to the inverse of the mean-reversion rate. Signal variability, quantified as the standard deviation after detrending, defined the noise amplitude. Numerical simulations were conducted using the Brian simulator with a time step aligned to the original data. Continuous wavelet transform was applied to the simulated OU series to decompose the signals into distinct frequency components.

### Retrograde labeling and immunostaining

Retrograde labeling, as previously described (57), was employed to define DA-WED neurons. Sharp quartz micropipettes (OD/ID: 1.2/0.6 mm, with internal filament) were pulled using a laser-based micropipette puller (P-2000, Sutter Instrument) to achieve tip resistances of 15–20 MΩ. Pipettes were backfilled with 5% Calcium Green-1 dextran (3,000 MW, anionic; C6765, ThermoFisher) in internal electrode solution, selected for retrograde transport. Iontophoretic injections were optimized to label neurons selectively while minimizing tissue damage, using voltage pulses (5–10 V, 500 ms, 1 Hz) delivered via an Analog Stimulus Isolator (Model 2200, A-M Systems) triggered by TTL pulses from a dual-channel waveform generator (DigiStim-2, ALA Scientific). Pulse patterns followed a GMM-distributed PSP profile characterized in this study. Injections lasted 10–15 minutes per site, with continuous monitoring of electrode resistance. The WED neuropil was visualized using IR-DIC optics with contrast enhancement (Olympus) and a Luigs & Neumann illumination system. Calcium Green fluorescence was detected using the PE4000 CoolLED illumination system (CoolLED Ltd., Andover, UK). Cell bodies displaying Calcium Green signal at expected locations (anterodorsal and lateral to the WED neuropil) were targeted for electrophysiological recordings. Following recordings, brains were fixed in 4% paraformaldehyde (2 h, 4°C), permeabilized with 0.3% Triton X-100, and immunostained for tyrosine hydroxylase (TH; 1:1000, AB152, Millipore) for 48 h at 4°C, followed by Alexa Fluor 594 secondary antibody (1:500, Invitrogen) and Alexa 488-conjugated streptavidin (1:100, E13345, Invitrogen) to visualize biocytin. Samples were washed in PBST several times at room temperature for 1 h, cleared in 70% glycerol in PBS for 5 min, and mounted in Vectashield (Vector Labs). Confocal images were acquired under 63× magnification using a Zeiss LSM800. Only neurons co-labeled with biocytin and TH were classified as verified DA-WED neurons, ensuring selective inclusion and excluding non-DA cells. Although Calcium Green and biocytin fluorescence spectra overlap, the Calcium Green signal was substantially fainter than biocytin at the confocal imaging stage, allowing reliable distinction between the two labels.

### Behavioral assay

Flies were yeast-deprived by maintaining them on 200 mM sucrose food (vials containing a Kimwipe soaked with 2 mL of sucrose solution). Flies were transferred to fresh sucrose food every other day. To achieve maximal protein starvation, males, virgin females, and mated females were deprived for 1-8 days, respectively. A blue dye stock solution was prepared by dissolving methylene blue (0.125 g per 10 mL distilled water). Test solutions were generated using the blue dye stock solution with yeast (1% w/v; 0.01 g yeast per 1 mL blue dye solution). Test vials were prepared per solution by layering the liquid feeding solution onto solidified agar. Vials were allowed to partially dry until mostly solid, and excess liquid was removed using a Kimwipe and disposable plastic tweezers immediately prior to use. Following the yeast-deprivation period, flies were transferred without anesthesia to test vials. Flies were allowed to feed for 30 min and were then immediately frozen at −20°F. After freezing, heads were removed and discarded using a razor blade. Fly bodies were collected in groups of 10 per sample, segregated by sex and subspecies, and transferred to Eppendorf tubes. Samples were homogenized in 300 μL PBS using a micropestle, followed by the addition of an additional 700 μL PBS. Homogenates were centrifuged at 13,000g for 20–25 min. Following centrifugation, 700 μL of the supernatant was transferred to clean tubes. Absorbance of the blue dye was measured at 625 nm using a NanoDrop spectrophotometer (Thermo Fisher Scientific). Fly locomotor activity and survival under starvation were monitored using Drosophila Activity Monitors (DAM; Trikinetics, Waltham, MA) in an incubator at 25°C under a 12 h:12 h light:dark (LD) cycle to quantify both starvation resistance and active periods.

## Statistical Methods

Data analyses were conducted using Prism version 10.6.1 (GraphPad), Clampfit version 10.7 (Molecular Devices), and MATLAB R2025b (MathWorks). For statistical analysis, two-group comparisons were conducted using unpaired t-tests. When comparing multiple groups, one-way ANOVA followed by Tukey’s post-hoc test was applied to normally distributed datasets, whereas the Kruskal-Wallis test with Dunn’s multiple comparisons test was used for non-normally distributed datasets. Differences were considered statistically significant at p < 0.05. Asterisks indicate significance levels: *p < 0.05, **p < 0.01, ***p < 0.001, and ****p < 0.0001. Error bars represent the standard error of the mean (SEM), averaged across all experiments.

## Acknowledgments

We thank Joseph Monaco for technical advice on computational modeling. We also thank Ben Strowbridge, Heather Broihier, and Dominique Durand, along with members of the Tabuchi lab for discussion, and the Light Microscopy Imaging Core at Case Western Reserve University for help with confocal microscopy. Funding: This work was supported by grants from the National Institutes of Health (R00NS101065 and R35GM142490), Whitehall Foundation, BrightFocus Foundation (A2021043S), PRESTO grant from Japan Science and Technology Agency (JPMJPR2386), Research Corporation for Science Advancement (SA-MBC-2024-080c), and the Tomizawa Jun-ichi and Keiko Fund of the Molecular Biology Society of Japan for Young Scientists.

---

**Fig. S1.** Additional characterizations of firing activity and intrinsic membrane properties in DA-WED neurons.

(**A**) Boltzmann sigmoidal model fitting analysis of time course plots for 1/interspike interval (ISI) of action potentials evoked by current injections. (**B**) Quantification of plateau arrival time duration based on fitting analysis of time course plots for 1/ISI of action potentials evoked by current injections. (**C**) Quantificaiton of half life time based on fitting analysis of time course plots for 1/ISI of action potentials evoked by current injections. during spontaneous firing. (**D**) Quantification of input resistance (GOhms) based on membrane voltage changes in response to current injections. Sample sizes: *D. melanogaster*: n=3, *D. melanogaster* with protein deprivation: n=3, *D. sechellia*: n=2, *D. sechellia* with protein deprivation: n=2. ns: non-significance based on unpaired t-tests. (**E**) Distribution of spike onset velocity (mV/s) values obtained from spontaneous action potential firing. Sample sizes (same data as in Fig. 1): *D. melanogaster*: n=7, *D. melanogaster* with protein deprivation: n=11, *D. sechellia*: n=4, *D. sechellia* with protein deprivation.

**Fig. S2.** Additional electrophysiological analysis of synaptic inputs in DA-WED neurons.

(**A**) Histogram showing inter PSP event intervals distribution in DA-WED neurons. (**B**) Histogram showing instantaneous frequency distribution in DA-WED neurons. Sample sizes: *D. melanogaster*: n=4, *D. melanogaster* with protein deprivation: n=7, *D. sechellia*: n=3, *D. sechellia* with protein deprivation: n=8. Sample sizes (same data as in Fig. 2 A-E): *D. melanogaster*: n=4, *D. melanogaster* with protein deprivation: n=7, *D. sechellia*: n=3, *D. sechellia* with protein deprivation: n=8. (**C**) Quantification of OU time constant (τ) defined as the exponential decay constant of the membrane voltage autocorrelation. (**D**) Quantification of OU noise amplitude (σ) defined as the variability of membrane voltage fluctuations. Sample sizes (same data as in Fig. 2 F-H): *D. melanogaster*: n=3, *D. melanogaster* with protein deprivation: n=6, *D. sechellia*: n=3, *D. sechellia* with protein deprivation: n=3. ns: non-significance based on unpaired t-tests.

**Fig. S3.** Additional electrophysiological analysis of DA-WED neurons.

(**A**) Histogram showing inter event interval distribution of spontaneous activity patterns of cardiomyocytes under control and yeast-deprived conditions in *D. melanogaster* and *D. sechellia*. (**B**) Quantification of inter event interval of spontaneous activity patterns of cardiomyocytes under control and yeast-deprived conditions in *D. melanogaster* and *D. sechellia*. Sample sizes (same data as in Fig. 3): *D. melanogaster*: n=8, *D. melanogaster* with protein deprivation: n=8, *D. sechellia*: n=8, *D. sechellia* with protein deprivation: n=4. ****p < 0.0001 based on one-way ANOVA followed by post-hoc Tukey tests.

## Notes

### Competing Interest Statement

The authors have declared no competing interest.

