## Supplementary figures and images for "Metaplastic neuronal state transition regulates species-specific interoceptive processing in *Drosophila*"

### Supplemental Data 1

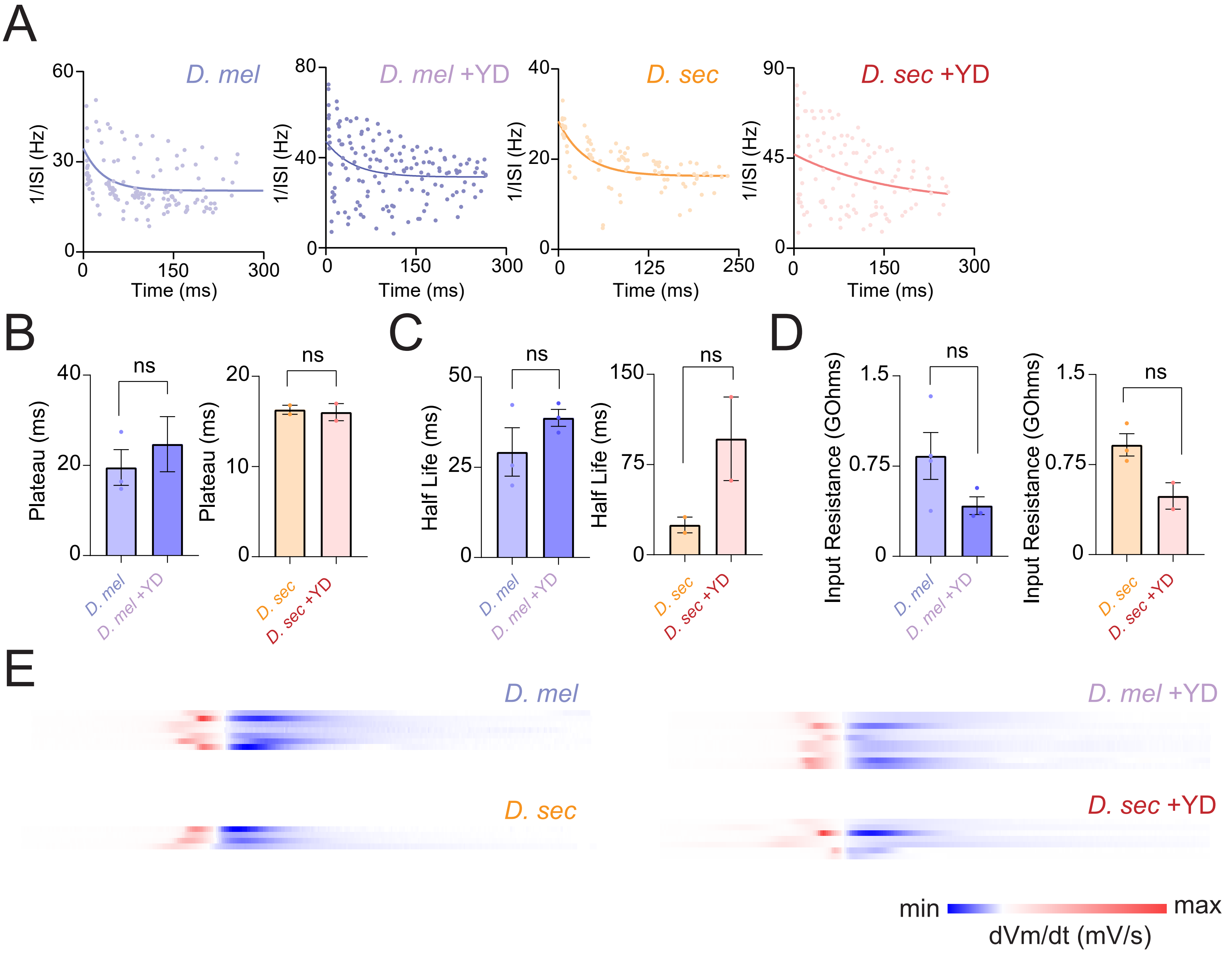

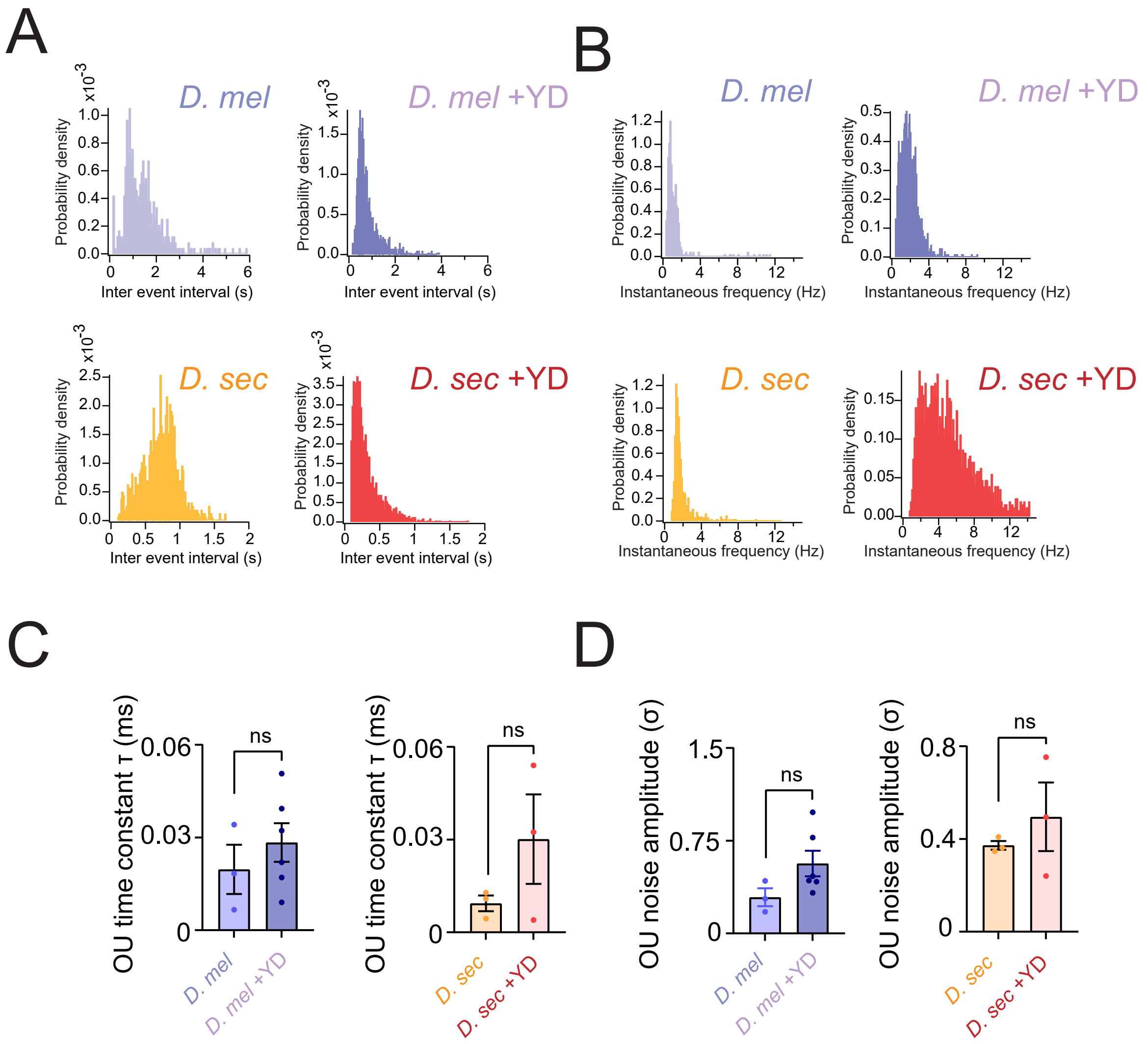

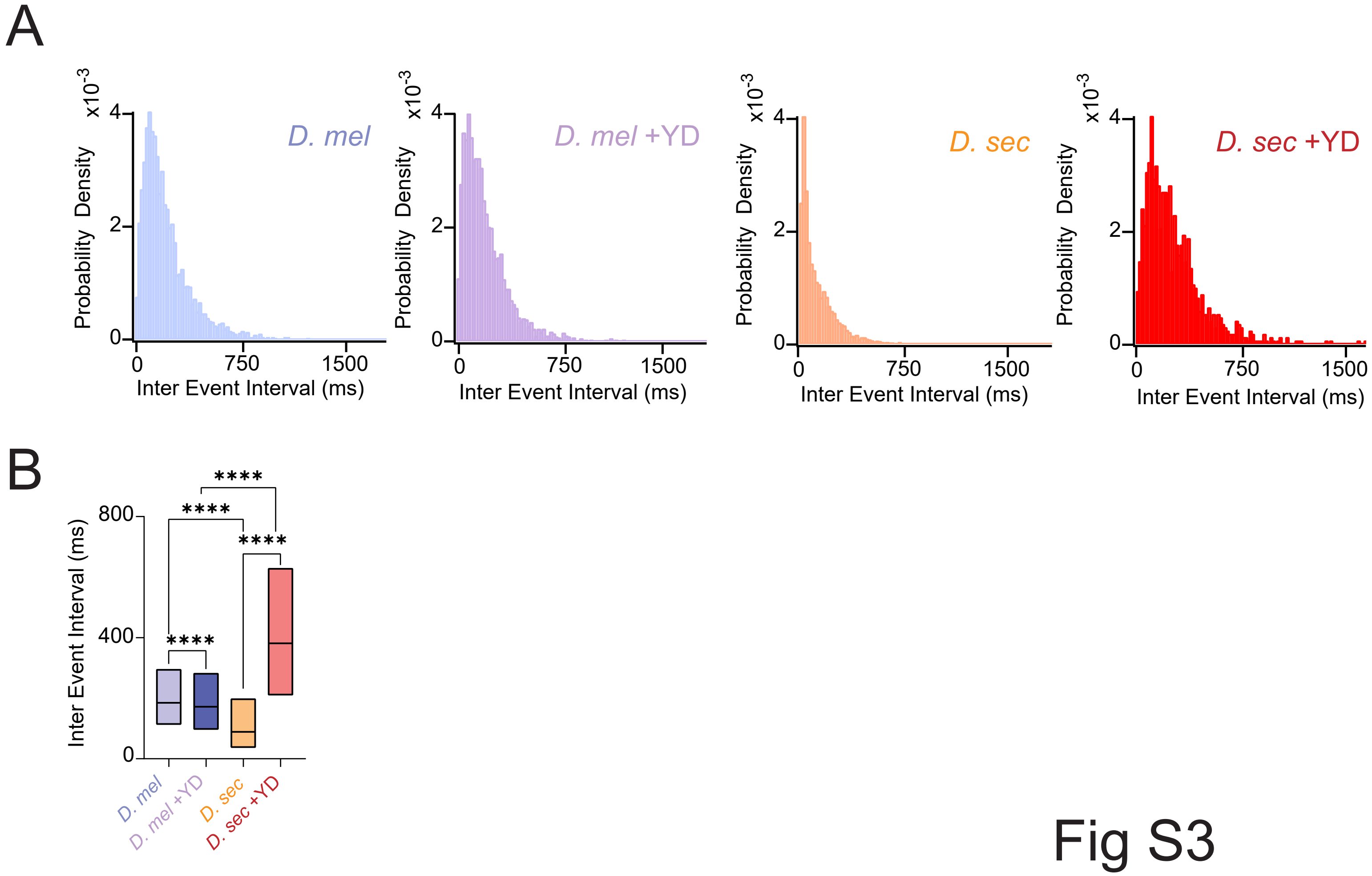
